# Can changes in ploidy drive the evolution to allogamy in a selfing species complex?

**DOI:** 10.1101/2023.11.27.568831

**Authors:** Ana García-Muñoz, Camilo Ferrón, Celia Vaca-Benito, María Nazaret Martínez-Gómez, Sílvia Castro, Mariana Castro, João Loureiro, A. Jesús Muñoz-Pajares, Mohamed Abdelaziz

## Abstract

- The evolution of mating systems in plants is central for understanding the rise of their diversity on Earth. The transition towards self-fertilization is a well-known example of convergent evolution although the opposite direction is expected to be forbidden according to evolutionary theories. We suggest that the ploidy level could promote changes in the reproductive strategies through its effect on traits related to pollination.
- We performed controlled crosses on several populations from the polyploid *Erysimum incanum* species complex, described as predominantly selfing, to evaluate the inbreeding depression. Additionally, we measured mating traits such as floral size, herkogamy, anther exertion, the relative investment in male and female components (P:O ratio) and genetic diversity.
- We described three ploidy levels in the complex – hexaploids were unknown until now. We found significant differences in the self-pollination success among ploidies and even among populations within the same ploidy. Inbreeding depression was present in higher ploidies, accompanied by bigger flowers with higher anther exposure, increased herkogamy and P:O and genetic diversity.
- These findings suggest that ploidy could be promoting alternative reproductive strategies to selfing, driving mating system diversification within a selfing species, which has not been previously described in the wild.

## Introduction

Patterns of genetic transmission between generations shape the mating system in every organism. Most flowering plants are hermaphrodites, developing pollen and ovules in the same flower. However, they display an impressive variation in mating strategies, including the development of organs and functions to optimise their mating success. The evolution of mating systems in plants and the mechanisms driving their diversity in nature have called the attention of scientists since Darwin (Darwin, 1877; Jain, 1976). Additionally, polyploidy is a pervasive phenomenon in flowering plants with significant implications on plant phenotype, physiology and biotic interactions (Oswald & Nuismer, 2011; Segraves & Thompson, 1999). Although increases in ploidy level have been widely associated with self-fertilisation for the stability of new karyotypes (Stebbins, 1950; Levin, 1975; Barringer, 2007), there is a lack of exhaustive studies exploring the mating system variation across more than two ploidies (Castro *et al*., 2013, 2020), especially when the ploidy variation is associated with an evolutionary transition to allogamy.

Although plants are sessile organisms, mating is not random since individuals have developed a wide variety of mechanisms to improve the effectiveness of the transmission of genes to the progeny. Reproductive strategies in plants range from self-compatible systems, promoted by autonomous or facilitated self-pollination, to self-incompatibility mechanisms, whose reproduction relies entirely on pollen transmission among individuals. Intermediate states between both strategies – known as mixed mating systems – are ubiquitous in the wild (Goodwillie *et al*., 2005). However, even though mixed mating is common (Aide, 1986; Barrett & Eckert, 1990), a transition towards selfing is expected to occur in most plant groups according to the evolutionary history of many taxa (Levin & Kerster, 1974; Zhang *et al*., 2022). The adaptive consequences of selfing are known since classical works from Darwin and Fisher, who hypothesised a high spreading ability of self-fertilisation alleles because of the transmission of the whole genetic material and the independence of mates and pollination vectors, conferring a reproductive assurance and granting colonising opportunities (Darwin, 1877; Fisher, 1949; Lloyd, 1979; Lloyd, 1992; Holsinger, 1996).

Mating system transitions involve relevant causes and consequences for genetic structures (Slotte *et al*., 2013; Novikova *et al*., 2023) and the evolutionary dynamics of populations (Allard, 1975; Charlesworth & Wright, 2001; Wright *et al*., 2013). However, speciation is expected to be reduced in selfing lineages, being considered evolutionary dead-ends (Takebayashi & Morrell, 2001; Igic & Busch, 2013). This hypothesis has been supported and well documented by many examples of convergent evolution to selfing, leading to a wide knowledge of the evolutionary consequences of this evolutionary transition (Stebbins, 1950; Grant, 1981; Charlesworth & Charlesworth, 1987; Charlesworth, 1992; Takebayashi & Morrell, 2001). However, the reverse transition – from selfing to outcrossing – has been considered unexpected to occurr in the wild and theoretically unfeasible (Takebayashi & Morrell, 2001; Igic & Busch, 2013). Even the long-term maintenance of mixed mating systems is controversial because they are considered an intermediate step in the transition from outcrossing to selfing (Flaxman, 2000; Plaistow *et al*., 2004; Goodwillie *et al*., 2005).

Selfing advantages are counteracted by fitness reductions experienced by inbred offspring, known as inbreeding depression (Darwin, 1877; Charlesworth & Charlesworth, 1987; Charlesworth & Willis, 2009). This reduction in reproduction success occurs when recessive deleterious alleles become unmasked as the frequency of homozygotes increases (Charlesworth & Charlesworth, 1987). In this scenario, the strength of inbreeding depression should evolve to reduce its effect in populations where self-pollination is frequent because deleterious alleles are purged by natural selection (Lande & Schemske, 1985; Schemske & Lande, 1985; Charlesworth & Charlesworth, 1987; Husband & Schemske, 1996). The limitation imposed by the low genetic diversity exhibited by selfing plants to cope with unpredictability in changing environments could be too constraining for the evolutionary outcomes of selfing populations. In contrast, cross-pollination in outcrossing species increases heterozygosity, which can be translated to more allelic combinations and even heterosis events. Predominantly outcrossing plants show inbreeding depression after self-pollination because the recessive deleterious alleles have not been purged. For this reason, the inbreeding depression index, which ranges from −1 to 1, is an accurate coefficient for characterising mating systems (Lohr & Haag, 2015).

Polyploidisation is the result of whole genomic duplications, having the potential to increase genetic diversity through new allelic combinations. Even though it is well-known how ploidy affects the plant phenotype (Jürgens *et al*., 2002; Te Beest *et al*., 2012; Moghe & Shiu, 2014), its influence on mating system evolution is a more arguing topic despite the high incidence of polyploidisation events across the evolutionary history of flowering plants (Grant, 1981; Soltis & Soltis, 1999; Otto & Whitton, 2000; Leebens-Mack *et al*., 2006; Soltis *et al*., 2009; Van de Peer *et al*., 2021). The effect of ploidy on mating system variation has been traditionally discussed from a theoretical framework which suggests that inbreeding depression is reduced in polyploid lineages, explaining the colonisation ability of polyploids in selfing species (Lande & Schemske, 1985; Barrett & Eckert, 1990; Husband & Schemske, 1996; Ronfort, 1999; Soltis & Soltis, 1999; Tate & Simpson, 2004; Barringer, 2007). However, only a few experimental works have compared populations differing in both ploidy and mating systems (Johnston & Schoen, 1996; Husband & Schemske, 1997; Miller & Venable, 2000; Rosquist, 2001; Mable, 2004), while studies addressing this question in populations sharing the same mating system are even scarcer (Barringer & Geber, 2008). Due to intra-specific variation in reproductive strategies and mating traits, considering several populations has gained importance for the accuracy of mating system characterization studies in recent years (Whitehead *et al*., 2018). Indeed, inbreeding depression estimation has been demonstrated to vary among populations (Barringer & Geber, 2008) and this variation could be more accentuated if such populations differ in ploidy level.

Different attributes are related to pollination and plant reproduction and, hence, shape organisms’ mating systems. Some involve primary sexual traits, such as reproductive investment or pollen-stigma recognition and rejection, while others are related to secondary sexual traits, such as flower attractiveness (Ornduff, 1969; Barrett & Harder, 1996). The corolla size might be one of the most representative traits related to plant attractiveness, enhancing pollinator visitation and pollen exportation (Herrera, 1996; Gómez *et al*., 2008; Caruso *et al*., 2019). In hermaphrodite plants, the relative performance of sexual organs plays an important role in pollination. While hermaphroditism can facilitate autonomous self-pollination in selfing plants, the co-occurrence of both sexual organs is problematic in plants exhibiting any significant level of inbreeding depression. Then, traits such as the anthers-stigma separation, known as herkogamy, are essential to avoid sexual conflict and subsequent problems derived from inbreeding depression (Charlesworth & Willis, 2009; Armbruster *et al*., 2014). Evaluating traits directly or indirectly related to reproduction is informative about the predominant reproductive strategy followed by plants. Genetic factors can influence the variation in these traits, which mediate the mating system evolution, as most of them have been demonstrated to have a genetic basis (Kruszewski & Galloway, 2006; Eckert *et al*., 2009; Karron & Mitchell, 2012; Cruzan & Barrett, 2016).

Regarding primary sexual traits and the sex allocation theory (Charnov, 1987, 1996), the relative relationships between pollen production and ovule amount per flower (i.e., P:O ratio) has been used as a conservative indicator in the mating systems characterisation (Cruden, 1977). This index is expected to show high values in outcrossing species since they must produce significant amounts of pollen which is subjetec to losses in the transportation pathway. Conversely, the P:O ratio is lower in selfing species because the arrival of pollen grains to the stigma of the same flower is assured. The values of the P:O ratio are often correlated with other floral traits, such as floral size and herkogamy (Galloni *et al*., 2007). Such correlations suggest that functions of all these traits are closely related between them (Stebbins *et al*., 1971; Barrett & Eckert, 1990; Lloyd *et al*., 1990) and with the plant mating strategy. The P:O ratio is important to recognise the mating system shown by a species, but it can vary among populations and even among individuals (Cruden, 2000). This is explained by the influence of environmental aspects on outcrossing and selfing rates, which are responsible for the mating system variation among locally adapted populations and even among years within the same population (Moeller & Gebre, 2005; Cheptou & Avendaño, 2006). Variation in the mating system involves relevant macro and microevolutionary consequences due to changes in population genetics and the interaction with other species (Charlesworth & Wright, 2001).

Here, we use the polyploid *Erysimum incanum* species complex, described as a selfing clade with most species being selfers (Nieto-Feliner & Clot, 1993; Fennane & Ibn-Tattou, 1999), to explore the role of polyploidy in shaping transitions in reproductive strategies from selfing to outcrossing. In particular, the main aims of this work were: (i) assess the ploidy levels of the studied populations of *E. incanum* species complex, (ii) characterise the mating system of each taxa and the main traits shaping them, (iii) explore the variation of secondary reproductive traits influencing the evolution of mating system in the species complex, and (iv) evaluate the consequences of mating system differentiation on the genomic variability within the group and its potential diversification. For that, traits closely related to the reproductive strategy, such as inbreeding depression, reproductive investment (and P:O ratio) and floral traits related to pollinator attraction were compared between different taxa of the *Erysimum incanum* species complex.

## Material and methods

### Study system

The genus *Erysimum* L. is one of the most diverse in the Brassicaceae, whose species are widespread throughout Eurasia, North and Central America and North Africa (Al-Shehbaz *et al*., 2006). Diversification in the genus was also promoted by patterns of local adaptation and hybridisation among lineages (Abdelaziz, 2013; Pajares, 2013). *Erysimum incanum* is considered a species complex, including annual and monocarpic species and subspecies inhabiting the Eastern part of the Iberian Peninsula, Southeast of France and the four main mountain ranges in Morocco (Nieto-Feliner *et al*. 1993; Fennane and Ibn-Tattou 1999; Abdelaziz *et al*. 2014). Diploid (2*n* = 2*x* = 16 chromosomes) and tetraploid (2*n* = 4*x* = 32 chromosomes; Nieto-Feliner & Clot, 1993; Fennane & Ibn-Tattou, 1999) populations are found within the species complex. Diploids of *E. incanum* present a vicariant distribution in the Rif and the Pyrenees mountains, where *E. incanum* subsp*. mairei* and *E. aurigeratum* were described, respectively. In the tetraploid level, we find *E. incanum* subsp*. incanum* that also presents a vicariant distribution in the Iberian Peninsula and Morocco (Nieto-Feliner & Clot, 1993; Fennane & Ibn-Tattou, 1999). *Erysimum incanum* subsp*. incanum* in Morocco was described as inhabiting the Middle Atlas, High Atlas and Antiatlas (Nieto-Feliner & Clot, 1993; Fennane & Ibn-Tattou, 1999). *Erysimum incanum* species complex exhibits a selfing syndrome, as the predominant phenotypes include small, hermaphroditic and self-compatible flowers (Feliner, 1991) and a recently described mechanism promoting prior selfing known as anther rubbing (Abdelaziz *et al*., 2019). Here, we studied nine populations of *E. incanum* whose geographical origin is detailed in Table 1.

**Table 1.** Summary of the populations (Pop.), their geographical origin and the number of individuals and crosses included in this study. The genetic information of each population is given by the genome size (G.S. mean ± SD) and the ploidy level (Ploidy).

| Ploidy | Pop. | Origin | N plants | N crosses | G.S. | ID | 99% C.I. |
| --- | --- | --- | --- | --- | --- | --- | --- |
| 2x | Ei12 | Rif | 67 | 196 | 0.35 ± 0.01 | -0.310 | **[-0.317,-0.302]** |
| 2x | Ei13 | Pyrenees | 19 | 34 | 0.34 ± 0.01 | -0.399 | **[-0.469,-0.331]** |
| 4x | Ei08 | Mid Atlas | 302 | 4013 | 0.78 ± 0.10 | -0.009 | **[-0.014,-0.005]** |
| 4x | Ei09 | Baetic | 110 | 572 | 0.78 ± 0.03 | -0.125 | [-0.162,0.088] |
| 4x | Ei11 | Baetic | 46 | 203 | 0.84 ± 0.07 | -0.111 | **[-0.121,-0.101]** |
| 6x | Ei16 | High Atlas | 90 | 527 | 1.10 ± 0.03 | 0.043 | **[0.017,0.070]** |
| 6x | Ei17 | High Atlas | 25 | 641 | 1.12 ± 0.04 | 0.562 | **[0.553,0.571]** |
| 6x | Ei18 | High Atlas | 74 | 543 | 1.13 ± 0.03 | 0.108 | **[0.098,0.118]** |
| 6x | Ei19 | High Atlas | 26 | 246 | 1.11 ± 0.02 | -0.390 | **[-0.397,-0.384]** |

### Genome size measurements

We collected young leaves to quantify the genome size using flow cytometry (FCM) and inferred the ploidy level following our previous knowledge of genome size variation in the *Erysimum* genus (Nieto-Feliner *et al*., 1993; Fennane *et al*., 1999; Muñoz-Pajares *et al*., 2018). We carried out the genome size analyses as described by Muñoz-Pajares *et al*. (2018). Between 10 and 20 individuals were analysed per population from every range where these species occur. Populations showing homogeneous values for ploidy level were considered to be composed mainly of a single cytotype. The ploidy levels found were studied as factors impacting reproductive strategies.

### Crossing experiment

Flowers from each population were subjected to two controlled hand pollination treatments: (1) Selfing (S), in which pollen grains from the own flower were deposited in the style, and (2) Intra-population Outcrossing (OC), in which flowers were pollinated with pollen from another individual from the same population. Flowers were carefully emasculated just before opening to avoid self-pollination before performing each treatment. At the end of the experiment, 6595 selfing and 380 outcrossing hand-made crosses were done on 759 *E. incanum* plants. The number of individuals and crosses in each population is detailed in Table 1.

### Crossing performance and inbreeding depression estimation

Fruits were collected to count the number of viable and aborted seeds and non-fertilised ovules. Then, three post-pollination fitness components were obtained and compared between *E. incanum* ploidy levels. ***Seedset*** is a fitness component that refers to the proportion of viable seeds produced per fruit compared to the total number of ovules, while ***fertility*** estimates the proportion of fertilised ovules (viable seeds and aborts) compared to the total number of ovules per fruit. A third fitness component, ***fertility success***, estimates the proportion of viable seeds compared to the number of fertilised ovules. The seedset for a treatment was considered zero when the fuit was not developed. These components are the result of pollen-stigma-ovule interactions which can fail at two different stages: (1) before fertilisation because of the inhibition of the pollen tube’s growth, which are not able to reach the ovary and the ovules are not fertilised, and (2) after fertilisation, when fertilized ovules fail to develop fully developed seeds. We visualised the pollen tube growth *in vivo* by UV microscopy following an analine blue staining protocol modified from Xie *et al*. (2017) by collecting self-pollinated flowers after 72 hours and preserving them in alcohol 90°.

The relative reproductive success of inbred line crosses compared to the outcrossing line is called ***inbreeding depression***. The inbreeding depression coefficient (ID) was calculated using only the seedset from selfing and outcrossing treatments by individual plant, population and ploidy following the Ågren & Schemske (1993):

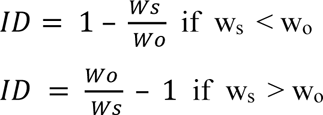

Where ***w_s_*** is the fitness component – the seed set in this case – from selfing treatment, and ***w_o_*** is the seedset from outcrossing treatment. Inbreeding depression ranges from −1 to 1. When it is negative and significantly different from zero, ***outbreeding depression*** occurs, meaning that selfing offspring show higher fitness than outcrossing ones (Frankham *et al*., 2011). Significant positive values correspond to ***inbreeding depression***, so fitness from outcrossing treatment is superior to selfing treatment. The significant values of these variables were calculated by computing the 99% confidence intervals using bootstrapping with 10,000 permutations, using the package *boot* v. 1.3-28 in R (Canty & Ripley, 2017). Values not significantly different from zero mean no fitness differences between selfing and outcrossing offspring.

### Mating traits measurement

For one flower in anthesis in a subset of 386 individuals, we measured the following phenotypic traits: corolla diameter (i.e., the distance between the edge of a petal and the edge of the opposite one), the length of the long stamens (i.e., the distance between the basis of the long filament and the anther) and the height of the style (i.e., the distance between the insertion point of the style in the basis of the corolla tube and the stigma surface). Short anthers were not considered as they do not contribute to spontaneous selfing. We calculated herkogamy as the difference between the stamen length and the style height, showing positive values when the stigma surface is above the stamens and negative values when the stigma is below the stamens, the latter facilitating the drop of own pollen on the stigma surface. We also estimated the anther exertion as the distance between anthers and corolla surface since this anther exposure potentilly influences pollen exporation.

Additionally, from these flowers, we collected half of the stamens (two long stamens and one short stamen) and preserved them in alcohol 70° for counting pollen grains. Male and female reproductive investments were estimated at the end of the plant life cycle as the overral pollen and ovule production per plant, respectively. We multiplied the number of pollen grains and ovules of a flower by the total number of flowers produced. Pollen grains and the number of ovules produced per flower were also used to calculate the P:O ratio described by Cruden (1977).

### Genetic diversity estimation

We obtained whole genome sequencing data from one individual per studied population. For that, we collected leaf material to be conserved in silica gel until DNA extraction, performed using the GenElute™ Plant Genomic DNA Kit (SIGMA) following manufacturer instructions. The purity, integrity and concentration of the resulting DNA were evaluated using agarose gel visualisations and spectrophotometric methods (Nanodrop and Qubit). Individual Illumina libraries were performed using the Collibri ES DNA Library Prep Kit (ThermoFisher) and sequenced by Novogene using a NovaSeq sequencing system. The resulting fastq files were aligned using bwa (Burrows-Wheeler Alignment Tool; Li & Durbin, 2009) and the *E. cheiranthoides* genome as a reference (http://erysimum.org/). We used samtools (Li *et al*., 2009; 2012) to convert sam files into bam files and bcftools (Danecek *et al*., 2021) to perform variant calls. Finally, the proportion of heterozygotic sites was estimated using VCFTools (Danecek *et al*., 2011).

### Statistical analyses

The autonomous selfing success and inbreeding depression were compared among ploidies and populations using ANOVA analyses and the Tukey test implemented in the R stat package. Also, the relationships among the corolla diameter, herkogamy, P:O ratio and inbreeding depression for different ploidies were performed using the package stat in R. We performed generalised linear mixed model analyses using the package lme4 v. 1.1.32 in R (Bates *et al*., 2009; Zuur *et al*., 2009) to explore the effect of treatments and ploidies (independently included as fixed factors and their interaction) on fitness components. These models were performed separately for seedset, fertility and fertility success. The individual plants nested within the population were included as random factors after performing caterpillar plots to visualise random effects. The different models were compared using package lme4 (Bates *et al*., 2009) and Akaike information criterion (AIC), Bayesian information criterion (BIC), log-likelihood (LogLik), and chi-squared test (*X^2^*). All analyses were performed using R Statistical Software (v4.2.1; Team, 2021).

## Results

### Genome size

We found three distinct and non-overlapping genome size ranges in the populations studied. According to our previous knowledge of genome size variation in the *Erysimum* genus (Nieto-Feliner & Clot, 1993; Fennane & Ibn-Tattou, 1999; Muñoz-Pajares *et al*., 2018), it was possible to assign each individual and population to a specific ploidy level, corresponding to diploids, tetraploids and hexaploids (Table 1). Diploid and tetraploid levels have previously been described in this species complex (Nieto-Feliner & Clot, 1993; Fennane & Ibn-Tattou, 1999), but the hexaploid cytotype is newly described in here. As the genome size was homogeneous in every population studied, we assumed that populations were composed of a single cytotype.

### Selfing and outcrossing success

We found significant differences in the reproductive success from selfing crosses among ploidies for all the estimated fitness components (Fig. 1a). Overall, seedset after selfing was higher in diploid populations and decreased with the increase in ploidy level (F = 89.0, *p*-value < 0.0001). We observed the same pattern for fertility (F = 108.1, *p*-value < 0.0001) and fertility success (F = 68.7, *p*-value < 0.0001). In diploids, fertility values were close to 1 indicating that all ovules were fertilised (Fig. 1a).

**Figure 1.**
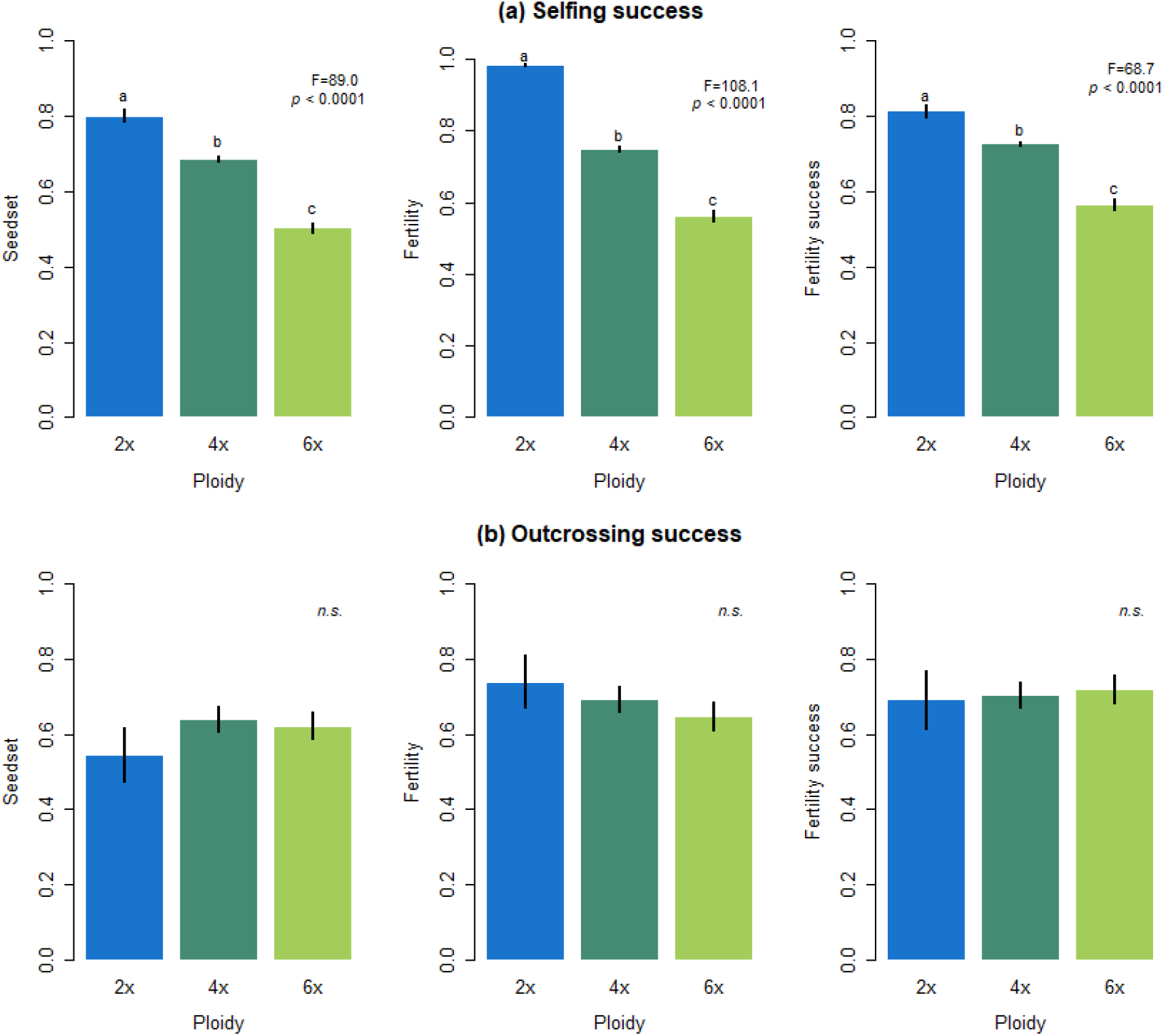
Mean values of fitness components estimated from (a) selfing and (b) outcrossing treatments for each ploidy. The fitness components were seedset, measured as the proportion of viable seeds by the total ovule production; fertility, as the proportion of fertilised ovules (seeds and aborts) by the total ovule production; and fertility success, as the proportion of viable seeds by the total number of fertilised ovules. Different letters indicate significant differences among ploidies according to the Tukey’s test. Significance *p*-values indicate the ANOVA results among ploidies (n.s = non-significant).

No significant differences were found among ploidies in any fitness components from the outcrossing treatment (Fig. 1b). GLMM analyses revealed that both pollination treatment (selfing or outcrossing) and the ploidy level (diploid, tetraploid or hexaploid) had an effect on the seedset, but the best explanatory model included the interaction between these two factors (Table 2). The analyses performed on fertility and fertility success showed the same pattern (Supporting Information Table S1 and S2).

**Table 2.** Outcome of the GLMM analyses testing the effect of the treatment and ploidy as fixed factors together with their interaction on *E. incanum* seedset. The individual plant and the population were considered as random factors nested within the ploidy level. Significant *p*- values are indicated in bold.

| Model | Seedset |  |  |  |  |  |
| --- | --- | --- | --- | --- | --- | --- |
| | AIC | BIC | logLik | $\chi^2$ | df | *p*-value |
| Intercept | 2973.4 | 2999.3 | -1482.7 |  |  |  |
| Seedset ~ Treatment + (1 \| Population/Plant) | 2970.8 | 3003.2 | -1480.4 | 4.539 | 1 | **<0.05\*** |
| Seedset ~ Ploidy + (Ploidy \| Population/Plant) | 2967.1 | 3070.8 | -1467.5 | 25.733 | 11 | **<0.01\*\*** |
| Seedset ~ Treatment * Ploidy + (Ploidy \| Population/Plant) | 2946.3 | 3609.4 | -1453.2 | 26.804 | 3 | **<0.0001\*\*\*\*** |

We found significant differences among populations in reproductive success from selfing crosses (F = 52.85; *p*-value < 0.0001), while such differences did not exist for the outcrossing treatment (Fig. 2). We observed that variation among populations increased in higher ploidies. This is, the two diploid populations did not differed significantly, while higher differences among populations, although non-significant, were observed in tetraploids, and accentuated differences were found among hexaploid populations. Surprisingly, we found the highest and the lowest values of seedset within hexaploids (Fig. 2a).

**Figure 2.**
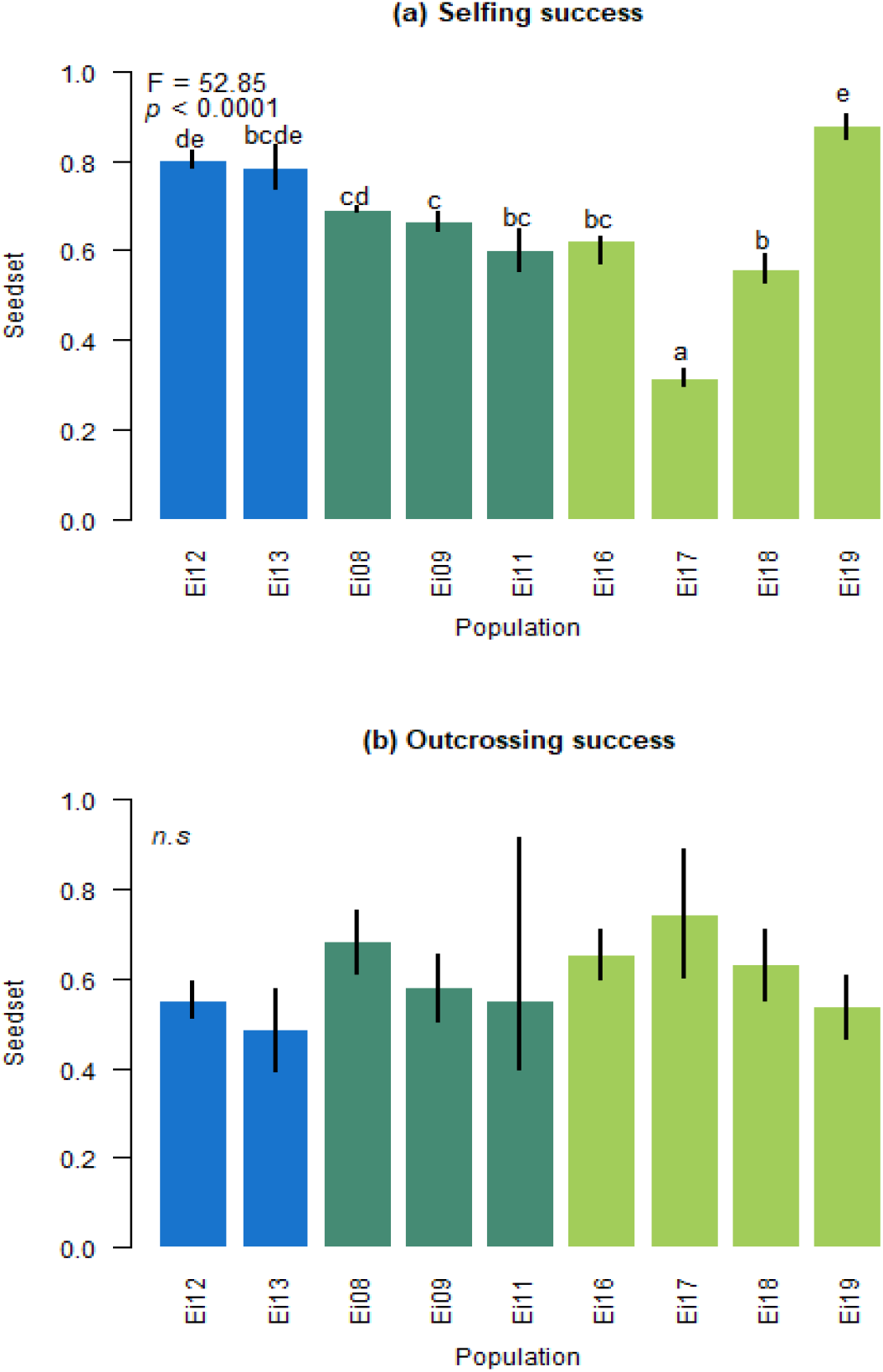
Mean values of seedset estimated from (a) selfing and (b) outcrossing treatments for each population. Blue bars refer to diploid, dark green bars to tetraploid, and green bars to hexaploid populations. Different letters indicate significant differences among ploidies according to the Tukey’s test. Significance *p*-values indicate the ANOVA results among ploidies (n.s = non-significant).

### Inbreeding depression

All populations showed significant values of inbreeding depression, except the tetraploid Ei09 population (Table 1). Diploid and tetraploid populations showed mostly significant negative values of inbreeding depression. Tetraploids showed less negative values of inbreeding depression than diploids, but still singifinact in 2 out of 3 populations (Ei08 and Ei11, Table 1). In contrast, significant and positive values of inbreeding depression were found in most hexaploid populations, except the Ei19 population where significant inbreeding depression values were obtained, similar to those exhibited by diploid populations (Table 1). However, when we compared the different ploidies (Fig. 3), they showed significant differences in inbreeding depression with increased values with increased ploidy. Thus, diploids showed a clear outbreeding depression pattern, while hexaploids showed the highest inbreeding depression values (F = 2257; *p*-value < 0.0001). Tetraploids showed values close to the non-difference between selfing and outcrossing performance (Fig. 3).

**Figure 3.**
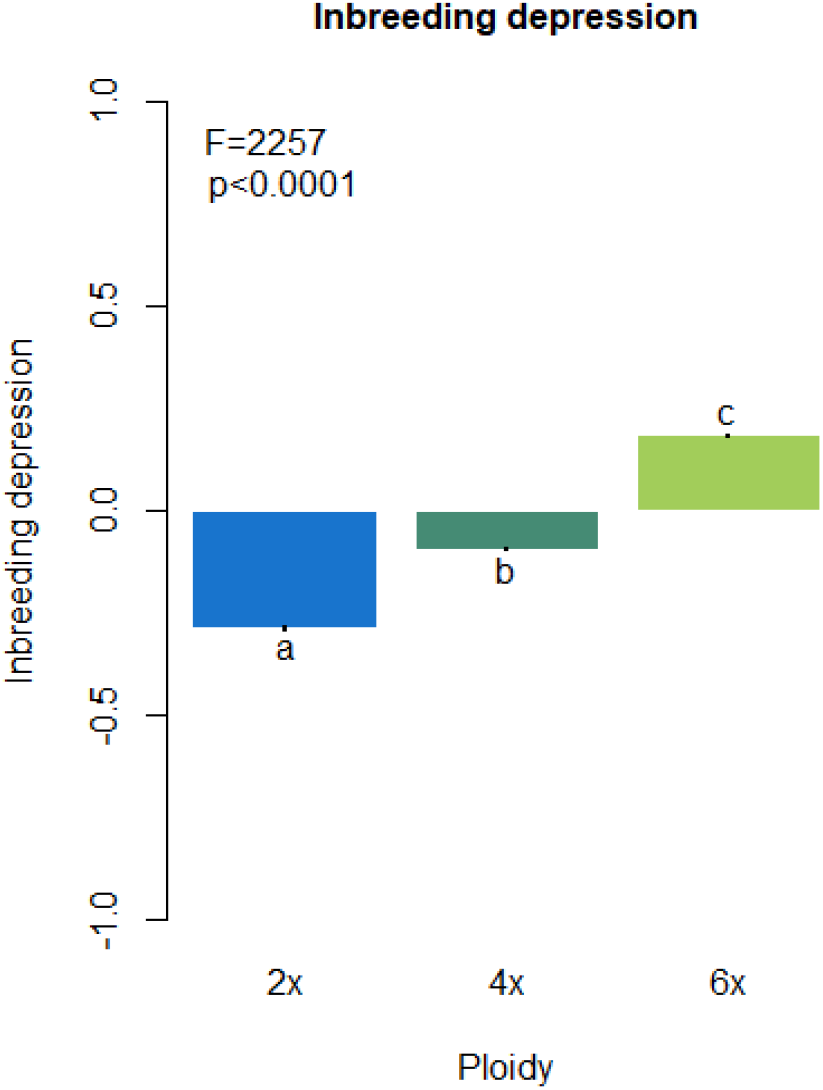
Mean values of inbreeding depression for each ploidy. Different letters indicate significant differences among ploidies according to the Tukey’s test. Significance *p*- values indicate the ANOVA results among ploidies.

### Reproductive investment

We found significant differences in male and female reproductive investment at the ploidy level. Hexaploids produced significantly more pollen than the other ploidies (F = 6.57, *p*-value < 0.01), while female reproductive investment was greater in tetraploids (F = 53.55, *p*-value < 0.0001). This was translated to a significantly higher P:O ratio in hexaploid plants (F = 18.35, *p*-value < 0.0001) (Fig. 4).

**Figure 4.**
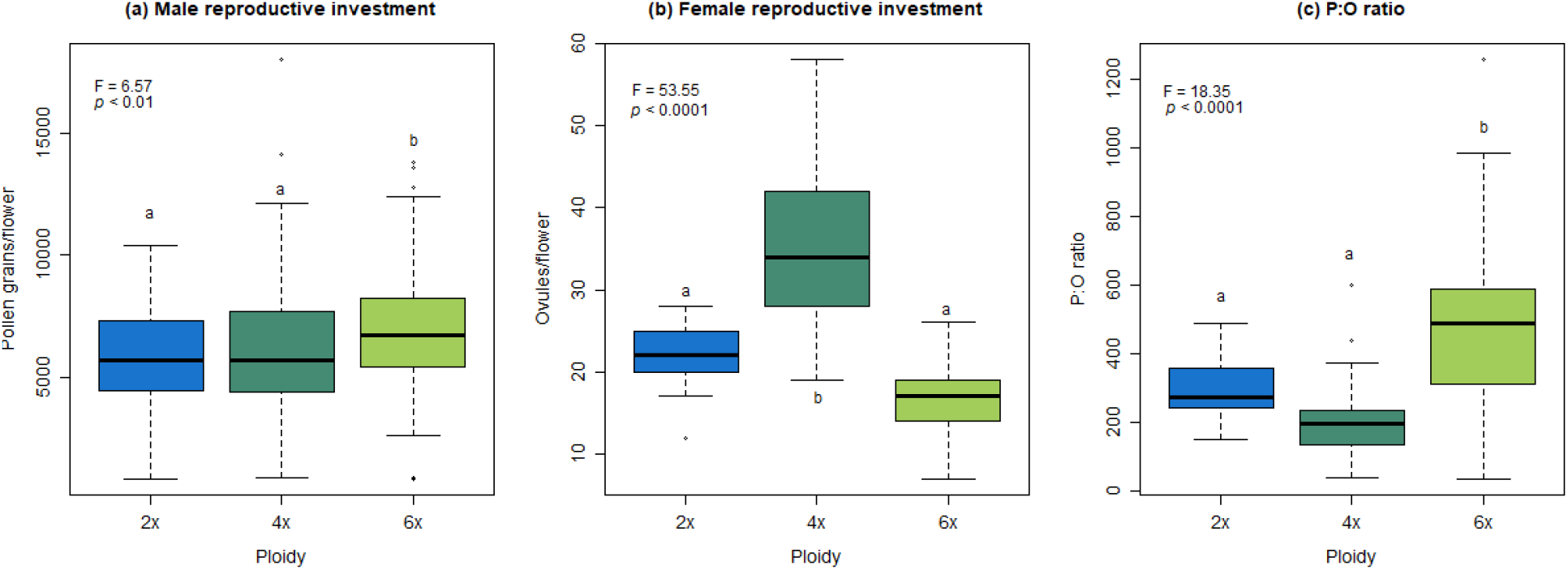
Reproductive investment among ploidies for (a) male function measured as pollen production, (b) female function measured as ovule amount and (c) the relative investment between male and female function estimated as P:O ratio. Different letters indicate significant differences among ploidies according to the Tukey’s test. Significance *p*-values indicate the ANOVA results among ploidies.

When we studied these variables at the population level, we found that hexaploid populations also produced fewer ovules (F = 15.51; *p*-value < 0.0001) and showed a higher P:O ratio than populations of other ploidies (F = 4.61; *p*-value < 0.001) (Fig. 5b,c). However, we found the most dispar values in pollen production in tetraploid populations, with Ei08 having the highest values and Ei11 the lowest pollen production values among all populations studied (F = 5.69, *p*-value < 0.0001) (Fig. 5a).

**Figure 5.**
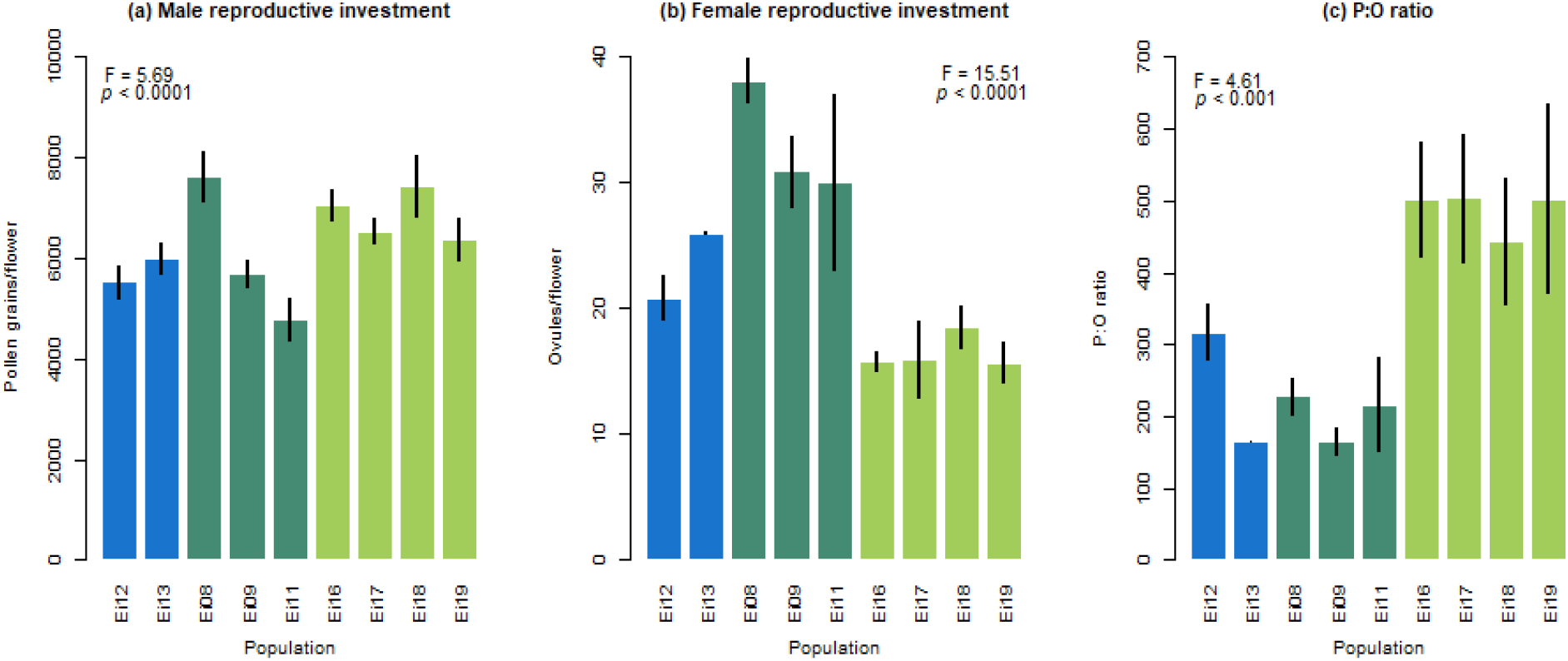
Reproductive investment among populations for (a) male function measured as pollen production, (b) female function measured as ovule amount and (c) the relative investment between male and female function estimated as P:O ratio. Blue bars refer to diploid, dark green bars to tetraploids and hexaploid bars to hexaploid populations. Significance *p*-values indicate the ANOVA results among ploidies.

### Mating traits

Significant differences were found in traits related to the mating system using a subset of 386 individuals from the three ploidies. Inbreeding depression was positive in hexaploids and negative in diploids (Fig 6a) as found above for the entire dataset of 759 individuals (Fig. 3). Tetraploids showed intermediate values of inbreeding depression between the two other ploidies and were not significantly different from zero (F = 3.66, *p*-value < 0.05) (Fig. 6a). The mean value of P:O ratio was higher in hexaploids (F = 18.35, *p*-value < 0.0001), while it was lower and not significantly different in diploids and tetraploids (Fig. 6b). Regarding floral traits, flowers were significantly bigger when ploidy increased (F = 51.63, *p*-value < 0.0001), with hexaploid plants showing the larger values of corolla size (Fig. 6c). Herkogamy was higher also in tetraploid and hexaploid plants (F = 12.71, *p*-value < 0.0001), indicating that polyploids showed a greater separation between sexual organs (Fig. 6d). We also found significant differences in the anther exertion values among ploidies (F = 31.25, *p*-value < 0.0001) (Fig. 6e) as the anther exposure was higher in polyploids.

**Figure 6.**
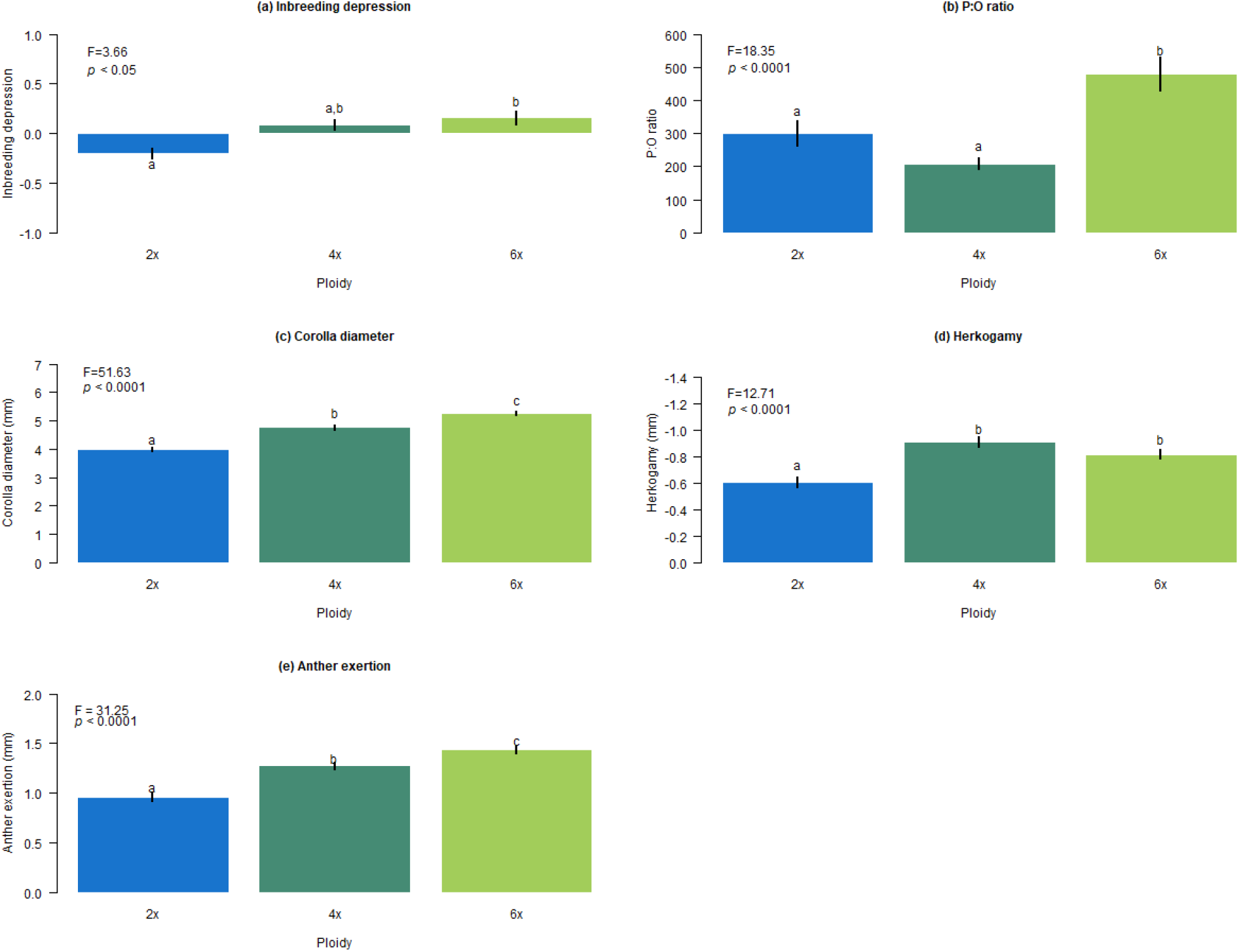
Mean values of traits related to the mating system: (a) inbreeding depression, (b) P:O ratio, and phenotypic traits: (c) corolla diameter, (d) herkogamy and (e) anther exertion measured in a subset of 386 plants. Different letters indicate significant differences among ploidies according to the Tukey’s test. Significance *p*-values indicate the ANOVA results among ploidies.

We found a significant and positive correlation between the P:O ratio and inbreeding depression in tetraploids (t = 2.44, df=26, *p*-value < 0.05), and this correlation was even stronger in hexaploids (t = 4.67; df = 17; *p*-value < 0.001) (Fig. 7a), i.e., plants less tolerant to their own pollen also produced proportionally more pollen compared to ovules. Additionally, plants with a higher P:O ratio showed larger flowers in hexaploids (t = 2.16, df = 21, *p*-value < 0.05) (Fig. 7b). Plants with a lower P:O ratio seemed to have lower herkogamy degree, especially in tetraploids; although this pattern was not significant (Fig. 7c). Inbreeding depression was not significantly correlated with corolla diameter for any ploidy (Fig. 7d). However, we found a significant positive correlation between inbreeding depression and herkogamy in diploids (t = 10.63, df = 3, *p-*value < 0.01) (Fig. 7e), i.e., plants with higher inbreeding depression showed less separation between sexual organs. Finally, we found that herkogamy was negatively correlated to corolla diameter in hexaploids (t= −3.56, df = 132, *p-* value < 0.001), indicating that larger flowers had a greater separation between sexual organs (Fig. 7f). Most of the significant relationships among mating traits are shown by the hexaploid level, with larger flowers producing more pollen in anthers more accessible to pollinators and more distant from the stigma, while showing higher levels of inbreeding depression.

**Figure 7.**
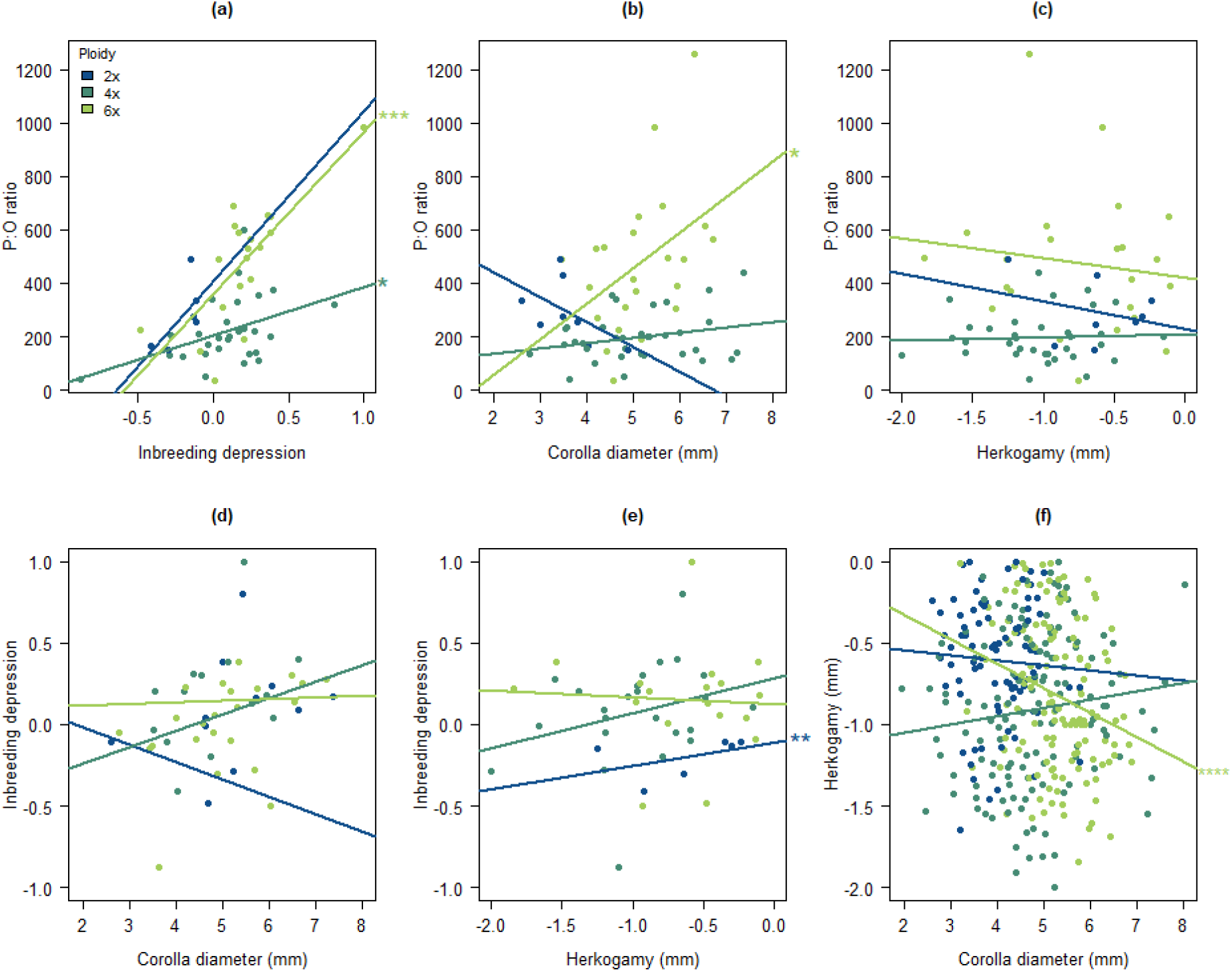
Correlation between pairs of mating traits: (a) P:O ratio and inbreeding depression, (b) P:O ratio and corolla size, (c) P:O ratio and herkogamy, (d) inbreeding depression and corolla size, (e) inbreeding depression and herkogamy and (f) herkogamy and corolla size. Asterisks indicate significant correlations. \**p*-value < 0.05; \*\**p*-value < 0.01; \*\*\**p*-value < 0.001; \*\*\*\**p*-value < 0.0001.

### Genomic diversity

We found a relevant difference in the level of heterozygosity between diploid and polyploid levels (F = 415.0, *p*-value < 0.0001) (Fig. 8). Genetic analyses revealed that polyploid plants have higher levels of heterozigocity than diploids. Interestingly, we found similar levels of genomic diversity in both tetraploid and hexaploid levels.

**Figure 8.**
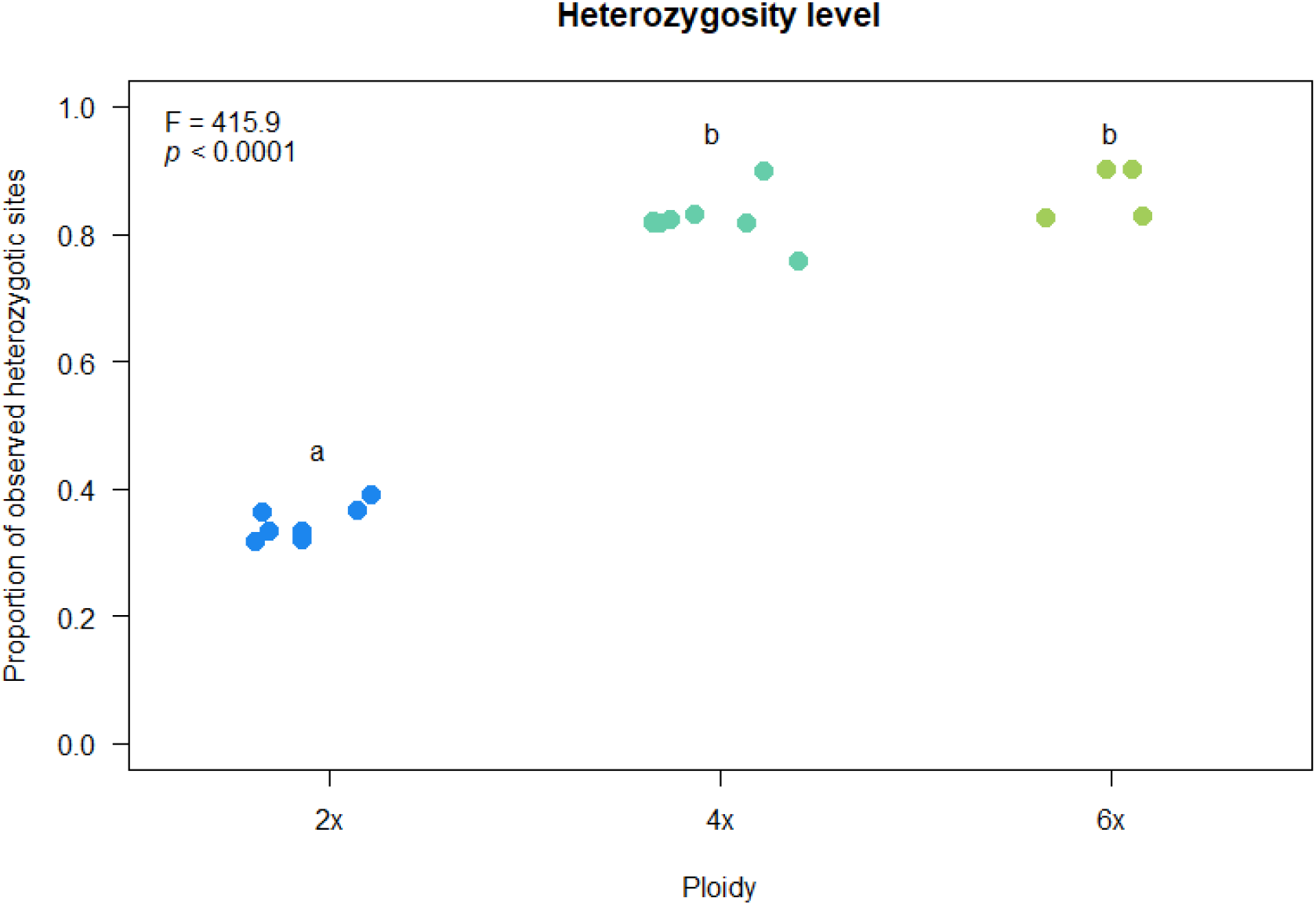
Heterozygosity levels of *E. incanum* ploidies. Different letters indicate significant differences among ploidies according to the Tukey’s test. Significance *p*- values indicated ANOVA results among ploidies (\**p*-value < 0.05; \*\**p*-value < 0.01; \*\*\**p*-value < 0.001; \*\*\*\**p*-value < 0.0001).

## Discussion

In this study, we explore the mating system variation of different populations of *E. incanum* and its association with ploidy changes and traits related to transitions between reproductive strategies. Regarding its reduced flower, pollen production, and levels of herkogamy, traits that facilitates self-pollination, and the overall lack of inbreeding depression, *E. incanum* is traditionally identified as selfing (Nieto-Feliner *et al*., 1993). Still, we found a gradient of tolerance to self-reproduction associated with the ploidy level – and even among populations within the same ploidy – varying from outbreeding to inbreeding depression and including intermediate values (Table 3).

**Table 3.** Summary of the traits used for characterising the mating system of the three ploidies found in *E. incanum*. Self. refers to the tolerance of individuals to their own pollen, I.D. refers to inbreeding depression, C.S. refers to corolla size, HK refers to herkogamy, and A.E. refers to anther exertion. Note that here were refer to the absolute magnitude of herkogamy (this is, higher separation between anthers and stigma) since its value in *E. incanum* is always negative.

| Ploidy | Mating traits |  |  |  |  |  |  | Predicted mating |
| --- | --- | --- | --- | --- | --- | --- | --- | --- |
|  | Self. | I.D. | P:O ratio | C.S. | HK | A.E. | Heterozygosity |  |
| 2x | ++ | -- | - | - | + | - | - | Selfing |
| 4x | + | - + | + | + | ++ | + | + | Selfing-Outcrossing |
| 6x | - | + | ++ | ++ | +++ | ++ | + | Selfing-Outcrossing |

Inbreeding depression is the main force opposing inbreed reproduction by optimising the reproductive success of outcrossing (Charlesworth & Charlesworth, 1987). However, this mechanism facilitates and even enhances the shift to inbreed reproduction after several selfing events through the purge of deleterious alleles (Lande & Schemske, 1985). Thus, the existence of inbreeding depression in a selfing species such as *E. incanum* is unexpected due to the expected lack of recessive deleterious alleles. In fact, we demonstrate the success of outcrossing events, which would potentially influence the population genetic structure, at least in hexaploids where positive inbreeding depression coefficients were found. The inbreeding depression found in polyploids was contrary to theoretical expectations (Lande & Schemske, 1985). In this sense, doubling gene copies after polyploidisation may have increased genetic diversity (just by chance, considering the mutation rate on bigger genomes) and may have a central role in masking recessive deleterious alleles, exhibiting more tolerance to selfing, at least during the first generations of self-reproduction (Siopa *et al*., 2020). By contrast, selfing tolerance is higher in diploid populations as early as the deleterious alleles are purged, a process that should be faster in lower ploidies. Indeed, the higher success of selfing compared to outcrossing treatments in diploid populations suggest that *E. incanum* shows outbreeding depression in its lower ploidies.

Beyond the ploidy, we also studied inbreeding depression at the population level and detected significant variation among populations. Previous studies have also showed tolerance to the own pollen, variation in inbreeding depression and outcrossing rates among different populations and years for the same species (Schoen, 1982; Lyons & Antonovics, 1991; Goodwillie & Ness, 2005; Vallejo-Marín *et al*., 2014), highlighting the importance of including among-population variation in characterising species mating systems (Whitehead *et al*., 2018). Population patterns observed within ploidies can result from different processes occurring at the intra-population level. The fact that mating systems can vary in space and time within a single species invites thinking about how mechanisms modifying self-fertilisation respond rapidly to natural selection (Jain, 1976). A deep understanding of such mechanisms would be interesting to explain the inter-population differences observed in *E. incanum*.

We tested inbreeding depression among using post-pollination fitness components as the consequence of pollen-stigma interaction. Both fertility and fertility success decreased with ploidy, butinterestingly, this pattern was more accentuated for fertility success. This result indicates that self-pollination failed immediately at the seed formation stage, while preliminary studies did not show differences in the growth of self-pollen tubes on stigma *in vivo* (Supporting Information Fig. S1). In addition, inbreeding depression can be estimated using different fitness components across offspring life, and the magnitude exhibited by each can be different (Jain, 1976). For example, classical works (Husband & Schemske, 1997) stated that inbreeding depression is more substantial in the earlier phases of offspring development, while other studies suggest that inbreeding depression varies its magnitude across post-dispersal stages of fitness (Grueber *et al*., 2010). Likewise, the rest of post-pollination fitness components could be interesting for inferring the consequences on genetic diversity and structure of the populations (Barrett & Harder, 2017) apart from testing inbreeding depression.

Ecological factors affecting outcrossing rates among populations are well described (Moeller & Gebre, 2005; Cheptou & Avendaño, 2006), but genetic factors seem less explored. Traits related to pollination could be affected by the ploidy level and our results show significant relationships among mating traits and ploidy, with hexaploid *E. incanum* having larger flowers, producing more pollen in anthers more accessible to pollinators and more distant from the stigma. The high P:O ratio and the overall high pollen production in hexaploids could indicate that a certain pollen amount can be exported instead of being used only for self-pollination. The P:O ratio has been demonstrated to help predict the reproductive strategy across several experimental works (Johnston & Schoen, 1996; Sato & Yahara, 1999; Fishman & Stratton, 2004; Goodwillie & Ness, 2005; Lozada-Gobilard *et al*., 2019) and remains a keystone in current pollination studies (Harder & Johnson, 2023). Interestingly, plants allocating more resources to pollen also show bigger flowers, which can serve to attract pollinators and promote pollen exportation (Etcheverry *et al*. 2012; Carleial *et al*. 2017). This positive relationship among mating traits might indicate that resources are jointly allocated to both primary and secondary sexual features and, thus, floral size and pollen amount are selected together.

Additionally, the levels of heterozygosity exhibited by polyploids support the occurrence of outcrossing events. The fact that both tetraploid and hexaploid levels showed similar heterozygosity levels suggests that this increase of heterozygosity compared to diploids is not due only to genome duplication. Instead, the pattern of genomic diversity observed in this study could be supported by the occurrence of outcrossing events in these populations. Recent studies in other polyploid species complexes have underlined the effects of the mating system and ploidy changes on genomic diversity (Lu *et al*., 2022). In our system, genomic diversity increased in higher ploidy levels. This result was relevant given that genetic diversity should be reduced in inbred populations, where evolutionary diversification is expected to be limited (Takebayashi & Morrell, 2001; Igic & Busch, 2013). We also demonstrated that ploidy potentially drove a shift to outbreeding in some polyploid populations. Therefore, the ploidy and mating system could jointly govern species diversification, as demonstrated in previous studies (Zenil-Ferguson *et al*., 2019). Additionally, although the origin of the polyploid populations (i.e., autopolyploid or allopolyploid) is still unknown, it would be key for better understanding the inter-population variation we found in polyploids.

Herkogamy is a key trait determining the levels of selfing and outcrossing (reference). Herkogamy together with flower size have been used as reliable indicators of the mating system, for example, in *Clarkia* species (Delesalle *et al*., 2008; Barringer & Geber, 2008), where species with small flowers showed reduced herkogamy – little stigma-anther separation – and were mainly self-pollinated while larger flowers showed a higher degree of herkogamy and were mainly outcrossing. These patterns were found in *E. incanum* in this study. The correlation between corolla size and herkogamy suggests that indirect selection would play a significant role in the evolution of these traits, which must be genetically correlated. Although flower size is more related to reproduction driven by pollinators, herkogamy avoids -or facilitates when it presents a low value-self-pollination. The role of herkogamy in avoiding sexual interference by hampering self-pollination is especially important in populations that show a strong inbreeding depression, such as hexaploid *E. incanum* populations. Therefore, a reproductive strategy different from self-pollination must be selected in hexaploids. The latter is well supported by our results of anther exertion. Polyploids exhibited a greater exposure of anthers above the corolla, enhancing the probability of pollen exportation by pollinators.

Genetic correlations between morphological and reproductive traits were found in previous studies (Stanton & Young, 1994; Young *et al*., 1994; Ashman, 2003), while the lack of such correlations in similar studies suggests that this covariance does not always exist (Mazer *et al*., 2007). Our results also revealed significant correlations between mating traits such as flower size and reproductive investment and the degree of self-pollen tolerance. The lack of significance in some of these correlations in hexaploids could be explained by the high variability in self-pollen tolerance since populations with contrasting values of inbreeding depression were found within the hexaploid level. The correlation between inbreeding depression and the P:O ratio was maintained for almost all ploidies, highlighting the accuracy of the P:O ratio for predicting the reproductive strategy (Cruden, 1977). In fact, Mazer *et al*. (2009) found that the P:O ratio was highly canalised in *Clarkia* selfing species while it was more variable in their outcrossing relatives. However, the P:O ratio increase seems to be driven by the inbreeding depression magnitude in *E. incanum*. In tetraploids, the correlation of inbreeding depression with corolla size might suggest that larger flowers are subjected to cross-pollination more than expected. A similar pattern was found in other *Erysimum* species (Abdelaziz *et al*., 2014a). An increase in flower size has a demonstrated effect on the frequency of visits by pollinators (Kennedy & Elle, 2008), which have an important role as selective pressures (Gómez *et al*., 2009) and as indicators of the outcrossing rate (Goodwillie *et al*., 2010). In our study system, we observe a change in corolla size among ploidies. The changes in corolla size would be the main force in modifying the pollinator attractiveness. We also found larger corollas with more herkogamy as ploidy increases. Previous studies described similar patterns (Webb & Lloyd, 1986; Barrett & Eckert, 1990), but opposite patterns were also described (Tate & Simpson, 2004). Thus, the ploidy variation can be associated with changes in the flower size but not necessarily with changes in selfing rate (Husband & Schemske, 1997). Although selfing and outcrossing rates were not strictly estimated in this study, the significant differences in several traits related to the type of pollination are relevant to infer changes in such wild population rates.

Our study found a wide variation of self-pollen tolerance among ploidies and even among populations within ploidies of *E. incanum* despite previously being described as predominantly self-compatible. Overall, self-compatibility is less intense when ploidy increases. In addition, self-pollen tolerance is accompanied by changes in other traits related to flower size, reproductive investment, and herkogamy. Diploids exhibit a proper selfing mating with significant outcrossing depression patterns, the smallest flower sizes and reduced herkogamy. However, hexaploid plants seem to experience significant inbreeding depression and exhibit higher values of flower size, herkogamy and P:O ratio. Our results highlight the importance of considering intraspecific variation to properly characterise the mating system of plant species and the potential role of ploidy to drive shifts in the mating system through changes in phenotypic traits playing a relevant role in reproduction. Patterns observed in this work may be especially relevant for shaping evolutionary trajectories among different populations and ploidies within a selfing species. Indeed, these changes mediated by the ploidy level may enhance reproductive strategies alternative to selfing, affecting population genetic structure, genome architecture and ecological interactions. The result of such processes could culminate in a transition toward outcrossing. The evolutionary transition in this direction has not yet been described and would be associated with speciation events.

## Acknowledgements

This article is dedicated to the beloved memory of María Jesús Ariza Molina (1982–2023), who sadly passed away too soon. We will always remember her everlasting smile and her warm help.

The authors thank Modesto Berbel, Celia Vaca-Benito, MariPaz Solis, Andrea Martín Salas, Cristobal Bragagnolo, Carlos Olmedo-Castellanos for lab assistance and paper discussion at the research network biochange (biochangenet.org). We also thank Luis Matías and María Jesús Ariza from the University of Sevilla for their helpful assistance and management to work in CITIUS. This research has been supported by a grant from the Spanish Ministry of Economy and Competitiveness (**CGL2014-59886-JIN**), the Organismo Autónomo de Parques Nacionales (Ref: **2415/2017**), and the Ministry of Science and Innovation (**PID2019-111294GB-I00/SRA/10.13039/501100011033**), including FEDER funds. AJM-P was funded by the European Commission under the Marie Sklodowska-Curie Action Cofund 2016 EU agreement 754446 and the UGR Research and Knowledge Transfer—**Athenea3i**. AG-M was supported by the *OUTevolution* project (**PID2019-111294GB- I00/SRA/10.13039/501100011033**). MC was supported by the Portuguese Foundation for Science and Technology (**SFRH/BD/89617/2012**).

## Competing interests

The authors disclose any conflicting competing interests.

## Author contributions

AGM, AJMP and MA thought and designed the experiments. AGM, CF, CVB and MNMG conducted the greenhouse experiments. MC, JL and SC conducted the ploidy analysis. AGM, CVB, AJMP and MA made the statistical analyses and designed the tables and figures. AJMP and AGM made the genomic analyses. AGM wrote the first draft of this manuscript, and the remaining authors made significant contributions to the draft. AJMP, SC, MC and MA got the funds to develop this study. AJMP and MA supervised the study.

## Data availability

Data and code is storage in https://drive.google.com/drive/u/0/folders/1I0b6PWgGriyFo3XtEh45qE5Gx6Yjcsc6 for review purposes, and will make it available in a public repository upon acceptance.

## Notes

### Competing Interest Statement

The authors have declared no competing interest.

