## Supporting Information for "Can changes in ploidy drive the evolution to allogamy in a selfing species complex?"

### *New Phytologist* Supporting Information

Article title: Can change in ploidy drive the evolution to allogamy in a selfing species complex?

Article acceptance date: Click here to enter a date.

The following Supporting Information is available for this article:

**Fig. S1** Visualisation of pollen tube growth on the stigma of a self-pollinated flower from diploid, tetraploid and hexaploid plants.

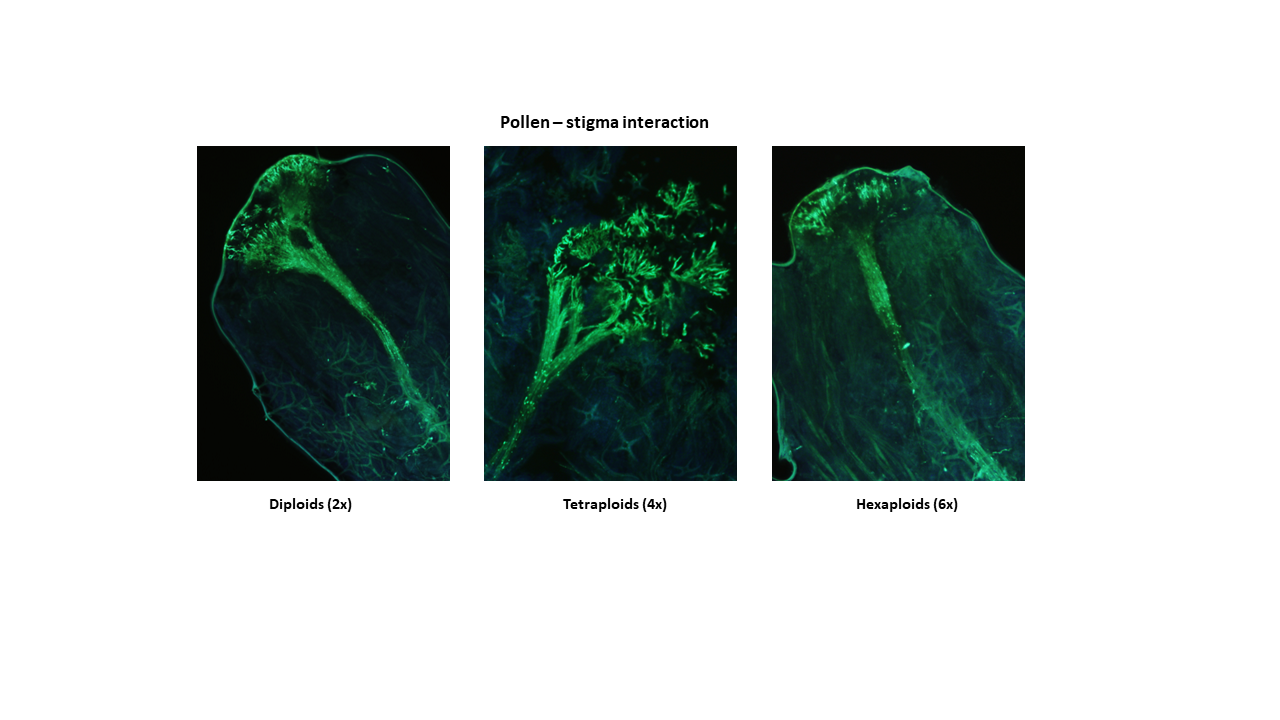

**Table S1** Outcome of the GLMM testing the effect of the treatment and the ploidy as fixed factors together with their interaction on *E. incanum* fertility. The individual plant and the population appear as random factors nested within the ploidy level. Significance *p*-values are indicated in bold (**p*-value < 0.05; ***p*-value < 0.01; ****p*-value < 0.001; *****p*-value < 0.0001).

|  | Fertility | | | | | |
| --- | --- | --- | --- | --- | --- | --- |
| Model | AIC | BIC | logLik | *χ^2^* | df | *p-*value |
| Intercept | 3411.4 | 3437.7 | -1701.7 |  |  |  |
| Fertility ~ Treatment + (1 \|Population/Plant) | 3399.4 | 3431.8 | -1694.7 | 13.977 | 1 | **<0.001***** |
| Fertility ~ Ploidy + (Ploidy \|Population/Plant) | 3399.2 | 3502.9 | -1683.6 | 22.186 | 11 | **<0.05*** |
| Fertility ~ Treatment * Ploidy + (Ploidy \|Population/Plant) | 2277.9 | 3501.0 | -1670.0 | 27.326 | 3 | **<0.0001****** |

**Table S2** Outcome of the GLMM testing the effect of the treatment and the ploidy as fixed factors together with their interaction on *E. incanum* fertility success. The individual plant and the population appear as random factors nested within the ploidy level. Significance *p*-values are indicated in bold (**p*-value < 0.05; ***p*-value < 0.01; ****p*-value < 0.001; *****p*-value < 0.0001).

|  | Fertility success | | | | | |
| --- | --- | --- | --- | --- | --- | --- |
| Model | AIC | BIC | logLik | *χ^2^* | df | *p-*value |
| Intercept | 3307.5 | 3326.8 | -1650.8 |  |  |  |
| Fertility success~ Treatment + (1 \|Population/Plant) | 3297.9 | 3323.8 | -1645.0 | 11.555 | 1 | **<0.0001****** |
| Fertility success ~ Ploidy + (Ploidy \|Population/Plant) | 3276.8 | 3341.4 | -1628.4 | 33.116 | 11 | **<0.0001****** |
| Fertility success ~ Treatment * Ploidy + (Ploidy \|Population/Plant) | 3257.6 | 3341.5 | -1615.8 | 25.261 | 3 | **<0.0001****** |
